# A cross-species interaction with a symbiotic commensal enables cell-density-dependent growth and in vivo virulence of an oral pathogen

**DOI:** 10.1101/2020.09.30.320770

**Authors:** Anilei Hoare, Hui Wang, Archana Meethil, Loreto Abusleme, Bo-Young Hong, Niki M. Moutsopoulos, Philip D. Marsh, George Hajishengallis, Patricia I. Diaz

**Affiliations:** Department of Oral Health and Diagnostic Sciences, School of Dental Medicine, UConn Health, Farmington, CT, 06030, USA; Laboratory of Oral Microbiology, Faculty of Dentistry, Universidad de Chile, Santiago, Chile; Department of Basic and Translational Sciences, Penn Dental Medicine, University of Pennsylvania, Philadelphia, PA, 19104, USA; Laboratory for Craniofacial Translational Research, Faculty of Dentistry, Universidad de Chile, Santiago, Chile; The Jackson Laboratory for Genomic Medicine, Farmington, CT, 06030, USA; Oral Immunity and Inflammation Unit, National Institute of Dental and Craniofacial Research, National Institutes of Health, Bethesda, MD, 20892, USA; Department of Oral Biology, School of Dentistry, University of Leeds, Leeds, UK; Department of Oral Biology, School of Dental Medicine, University at Buffalo, State University of New York, Buffalo, NY, 14215, USA

**Keywords:** inter-species interactions, cooperation, cell-density-dependent cues, microbial succession, microbial communities, pathogen colonization and virulence

## Abstract

Recent studies describe in detail the shifts in composition of human-associated polymicrobial communities from health to disease. However, the specific processes that drive the colonization and overgrowth of pathogens within these communities remain incompletely understood. We used in vitro culture systems and a disease-relevant mouse model to show that population size, which determines the availability of an endogenous diffusible small molecule, limits the growth, colonization, and in vivo virulence of the human oral pathogen *Porphyromonas gingivalis*. This bacterial pathogen overcomes the requirement for an endogenous cue by utilizing a cell-density dependent, growth-promoting, soluble molecule provided by the symbiotic early colonizer *Veillonella parvula*, but not produced by other commensals tested. Our work shows that exchange of cell-density-dependent diffusible cues between specific early and late colonizing species in a polymicrobial community drives microbial successions, pathogen colonization and disease development, representing a target process for manipulation of the microbiome towards the healthy state.

## Introduction

Diffusible signals allow bacteria to coordinate behaviors such as bioluminescence, competence, biofilm formation, sporulation and virulence, according to the size of the population (*1*). A less studied form of cell-to-cell communication is that which is required for replication. In several bacterial species, the size of the inoculum is a critical determinant of in vitro growth (*2*). Such an inability to grow at low cell-density is relieved by addition of conditioned spent medium from the same species, highlighting the endogenous nature of the required cue. A dependency on autoinducing molecules to grow is likely to limit colonization of new habitats by bacterial populations at a low cell-density. Bacteria could overcome this requirement by establishing in pre-existing polymicrobial communities, where resident species provide the growth-initiating cues that newcomers need.

The human oral cavity, in particular teeth and the gingival sulci, harbor diverse microbial communities. These polymicrobial biofilms represent an accessible model in which to study the role of inter-species interactions in community assembly and development processes. The compositional shifts during oral community maturation have been described in detail (*3–5*), with early colonizers creating niches conducive to the establishment of later and often anaerobic colonizers. If oral hygiene fails to restrict biomass accumulation and species successions continue, an inflammatory response in the adjacent gingiva is triggered and is referred to as gingivitis (*4*). In some individuals, communities undergo further compositional shifts resulting in overgrowth of even more pathogenic species, which trigger periodontitis, an inflammation-mediated destruction of tooth-supporting tissues that leads to tooth loss and constitutes a risk factor for several systemic diseases (*6*). Early and late oral biofilm colonizers have been shown to cooperatively interact to degrade host macromolecules, to establish reduced (i.e. anaerobic) environments and to exchange metabolic byproducts, thereby driving community maturation and subverting host defenses (*7–9*). However, the role of population-dependent inter-species communication on microbial successions and the emergence of dysbiosis remains unclear. Whether late colonizers require growth-initiating factors that otherwise limit their establishment during early biofilm development has not been investigated.

*Porphyromonas gingivalis*, an anaerobic late-colonizer, becomes an abundant species in dental communities of subjects affected by periodontitis (*10*). *P. gingivalis* has been associated with progression of human periodontitis, and shown to dysregulate immune surveillance leading to bone loss, the hallmark of periodontitis, in animal models (*11–13*). *P. gingivalis* has difficulty in becoming established in the oral cavity as shown by its presence as a transient commensal in children and its low abundance, when present, in early dental biofilms (*4, 14, 15*). While a reduced atmosphere created by early colonizers and the availability of inflammation-derived proteinaceous nutritional substrates are probably required for the establishment of *P. gingivalis* in the gingival crevices (*16*), an inability to grow at low cell-density might also contribute to late colonization by this species. Routine laboratory growth of *P. gingivalis*, especially in chemically-defined medium, requires a large inoculum (*17*). Accordingly, we investigated whether *P. gingivalis* requires a cell-density-dependent autoinducing signal to grow and whether this cue could be provided by early biofilm colonizers. We present evidence that the growth of *P. gingivalis* is controlled by a diffusible cell-density-dependent small molecule. Such a dependency on an autoinducer is overcome by an inter-species interaction with the early colonizing commensal *Veillonella parvula*, which allows low-cell-density *P. gingivalis* to grow in vitro and also to colonize the mouse oral cavity, where it promotes periodontal bone loss. Our work shows that although growth of the oral pathogen *P. gingivalis* depends on an autoinducing diffusible small molecule, a cross-species interaction with an early colonizing symbiotic commensal enables pathogen colonization and virulence.

## Results

### Growth of *P. gingivalis* is dependent on a soluble factor produced at high cell-density

The growth of *P. gingivalis* in a nutrient-restricted medium supplemented with an iron source and host macromolecules (mucin-serum) was found to be dependent on the initial cell-density as batch cultures inoculated with less than 10^7^ cells mL^−1^ were unable to grow (Figure 1a). Identical inoculum size thresholds were seen for three different strains of *P. gingivalis* (Figure 1a and Supplemental Figures 1a and 1b). However, growth from a low-cell-density inoculum (10^5^ cells mL^−1^) was possible in the presence of cell-free spent medium from a *P. gingivalis* early stationary phase culture, suggesting that growth initiation was dependent on an endogenous soluble factor that had accumulated in the medium (Figure 1b and Supplemental Figure 1c). Remarkably, even 100% unsupplemented spent medium was able to support growth, with these cultures reaching comparable maximum densities to those grown in the presence of fresh medium.

**Figure 1.**
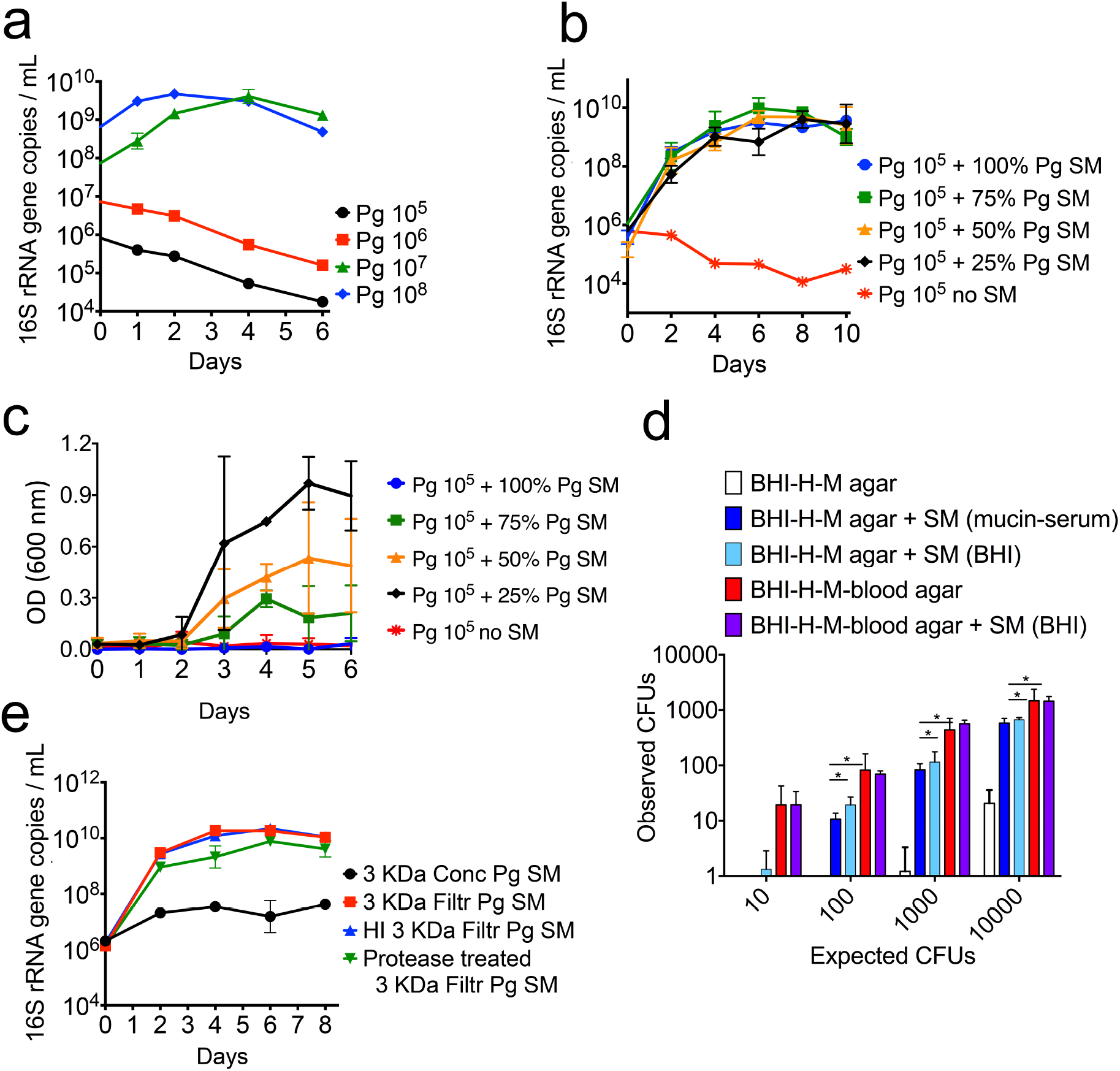
Growth of *P. gingivalis* (Pg) is dependent on a soluble factor accumulated at high cell-density. **a.** Growth of Pg strain 381 in mucin-serum liquid medium is dependent on the size (cells mL^−1^) of the inoculum. Cultures inoculated at various cell densities were incubated and sampled anaerobically, followed by determination of growth via qPCR. **b.** Mucin-serum spent-medium (SM) from a Pg 381 stationary phase culture restored the growth of a low-cell-density inoculum (10^5^ cells mL^−1^) of Pg 381. Pg was inoculated in fresh mucin-serum containing the indicated proportion (vol/vol) of SM. **c.** SM from a Pg 381 stationary phase culture grown in brain heart infusion (BHI) supplemented with cysteine, hemin (H) and menadione (M) restored the growth of a low-cell-density inoculum (10^5^ cells mL^−1^) of Pg 381. Pg was inoculated in fresh BHI-H-M containing the indicated proportion (vol/vol) of SM and growth was monitored via optical density (OD). **d.** SM from Pg 381 augmented the number of colony forming units (CFUs) recovered on BHI-H-M agar. Pg was diluted in PBS or SM and plated at different densities. The number of CFUs obtained was compared to the number expected according to microscopic counts. * represents a p value <0.05 as determined by t tests. **e.** Pg soluble factor capable of supporting its growth from a low-cell-density inoculum is smaller than 3 kDa, is heat-stable and is protease resistant. SM from Pg 381 grown in mucin-serum was filtered through 3 kDa membranes and either heat-inactivated (HI) or treated with proteases, followed by lyophilization and reconstitution (10x) in dIH20. Reconstituted fractions (Conc = > 3kDa and Filtr = < 3 kDa) were added to fresh mucin-serum medium (1:3, vol:vol) to evaluate growth of low-cell-density (10^5^ cells mL^−1^) Pg. Data in all panels represent replicates (mean and standard deviation) from at least three independent experiments.

A 10^5^ cells mL^−1^ inoculum also failed to grow in a different medium (BHI-H-M), but again spent medium from a stationary-phase culture restored growth (Figure 1c). In BHI-H-M, however, growth in the presence of spent media was less consistent across replicates (n=6) and higher proportions of fresh medium supported higher maximum densities. The effect of spent medium was also tested on solid BHI-H-M, where resuspension of the inoculum in spent medium from stationary-phase liquid cultures, grown either in mucin-serum or BHI-H-M, significantly increased the number of colony forming units (CFUs) recovered (Figure 1d). The addition of blood, which is commonly incorporated into solid media to grow *P. gingivalis*, allowed the number of observed CFUs to approximate the expected level (based on microscopic counts). In the presence of blood, spent media did not further augment the number of recovered CFUs (Figure 1d).

To characterize the nature of the soluble factor(s) that facilitated growth of a low cell-density inoculum of *P. gingivalis*, the spent medium of a stationary phase culture grown in mucin-serum was fragmented with a 3kDa-MWCO filter, and both the concentrate and filtrate (fraction <3kDa) were tested for activity. Only the filtrate supported growth of *P. gingivalis* (Figure 1e). Filtrates of a 1kDa-MWCO membrane also enabled growth (Supplemental Figure 1d). Furthermore, the growth-promoting activity of filtrates was heat-stable and protease-resistant (Figure 1e).

Altogether, these data show that growth of *P. gingivalis* requires a threshold concentration of a soluble endogenous heat-stable and protease-resistant small molecule. Growth can only occur when cells are transferred to fresh medium at a density that allows accumulation of the molecule to occur or in the presence of spent media containing the growth-promoting factor.

### Known quorum-sensing mediators do not support growth of low cell-density *P. gingivalis*

A set of compounds previously found to mediate inter-cellular communication were tested for their ability to stimulate growth of a low-cell-density inoculum of *P. gingivalis* (Supplementary Table 1). Supplementation of mucin-serum medium with D-pantothenic acid (D-PA), which regulates growth of low-cell-density *Cryptococcus neoformans* (*18*), or with the metabolically-related molecules panthenol and β-alanine, had no effect. Tyrosol, a quorum-sensing molecule that supports growth of low-cell-density cultures of *Candida albicans* (*19*), also failed to stimulate *P. gingivalis*. A set of polyamines, including spermidine, spermine, cadaverine and putrescine, which stimulate eukaryotic and prokaryotic cell growth (*20*) had no effect. The addition of 4-aminobenzoate/para-amino benzoic acid (pABA), which is needed for maximal biofilm accumulation of *P. gingivalis* (*7*), also failed to stimulate growth. The LuxS system was not involved, since a *P. gingivalis* Δ*luxS*::*ermF* mutant (*21*) showed similar behavior to the wild-type strain, only growing in mucin-serum when inoculated at high cell-density; and spent medium from the Δ*luxS* strain supported growth of a low-cell-density inoculum of wild-type *P. gingivalis* (Supplemental Figure 1e).

### Early colonizers do not exhibit cell-density-dependent growth and enable growth of low-cell-density *P. gingivalis*

In a cross-sectional evaluation of publicly available 16S rRNA gene datasets from human subjects with periodontal health, gingivitis and periodontitis, it is clear that *P. gingivalis* exhibits a progressively higher frequency of detection and abundance in subgingival biofilms as periodontal health deteriorates (Figures 2a and 2b). We next evaluated if species present during early stages of biofilm dysbiosis could support the growth of low-cell-density cultures of *P. gingivalis.* We tested five species with high prevalence and abundance in gingivitis (Figures 2c and 2d). Some of these species were also present in high proportions in health, but we reasoned that since the total microbial load increases by at least 3-log from health to gingivitis (*4*), these prevalent species and the diffusible molecules they produce would accumulate during the gingivitis state to a threshold that may allow the establishment and growth of *P. gingivalis*. As seen in Figure 2e, co-inoculation of *P. gingivalis* in mucin-serum with the five early colonizing microorganisms facilitated its growth from even a low-cell-density inoculum. The early colonizers all grew within this community reaching their maximum yield in 2 days, while *P. gingivalis* reached a biomass after 6 days that was comparable to that achieved when inoculated alone at high cell-density (as shown in Figure 1a).

**Figure 2.**
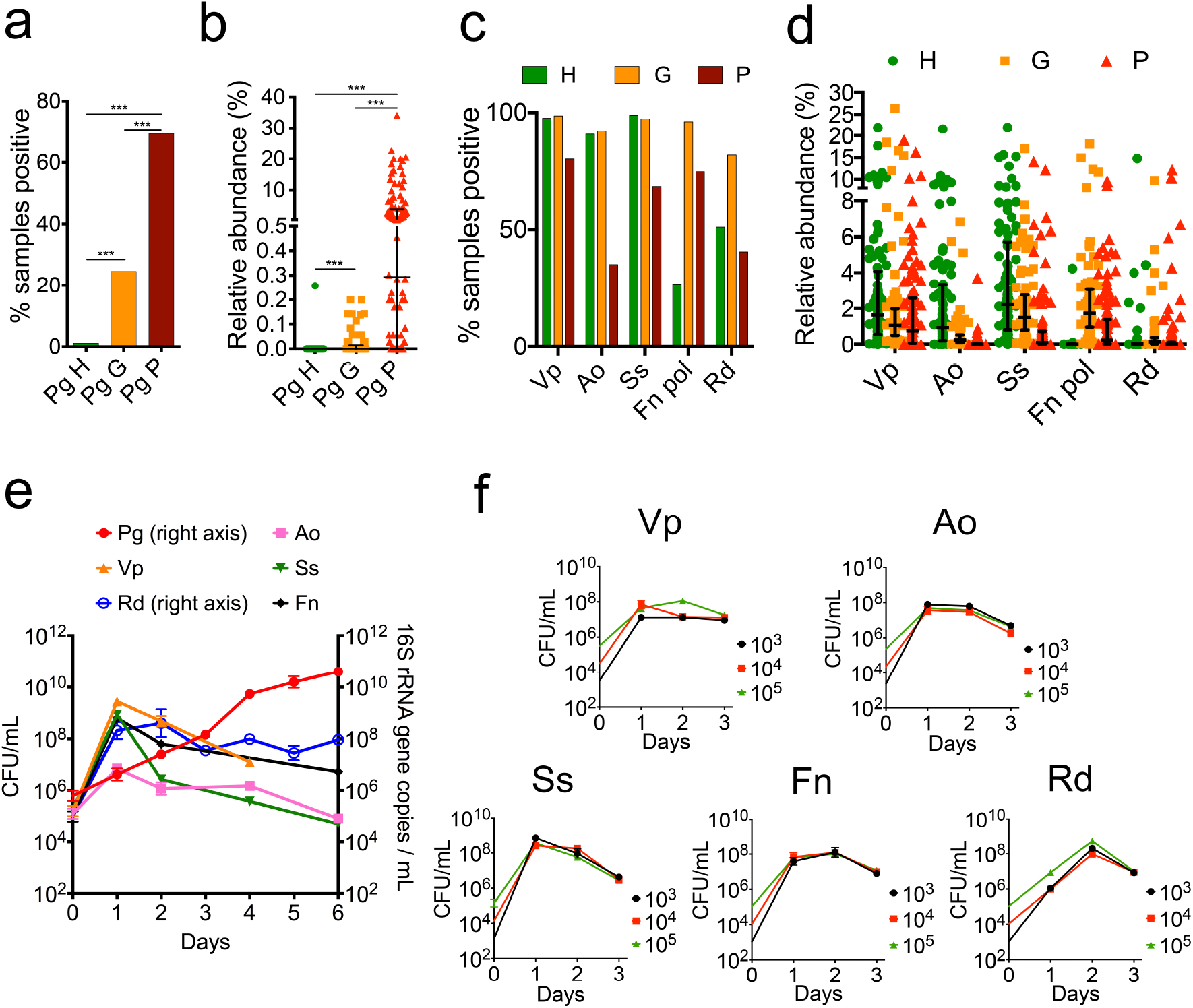
Growth of low-cell-density *P. gingivalis* (Pg) is supported by a community of species that are abundant in early subgingival biofilms. Detection (**a**) and relative abundance (**b**) of Pg in subgingival plaque in states of periodontal health (H), gingivitis (G) and periodontitis (P). Detection (**c**) and relative abundance (**d**) of *Veillonella parvula* (Vp), *Actinomyces oris* (Ao), *Streptococcus sanguinis* (Ss), *Fusobacterium nucleatum* subsp. *polymorphum* (Fn pol) and *Rothia dentocariosa* (Rd) in subgingival plaque at different disease stages. Lines in relative abundance graphs represent median and interquartile range. **e.** Co-inoculation of Pg with Vp, Ao, Ss, Fn and Rd in mucin-serum results in growth of all species and enables growth of a low-cell-density inoculum of Pg. All species were inoculated together, each at a density of 10^5^ cells mL^−1^ and cultures incubated and sampled under anaerobic conditions. Cell numbers of Vp, Ao, Ss and Fn were determined by plating on selective media. Biomass of Pg and Rd was determined via qPCR. **f.** Individual growth of species in the supporting community is not cell-density-dependent as shown by the ability of all species to grow in monoculture in mucin-serum when inoculated at a density as low as 10^3^ cells mL^−1^.

We next tested whether the early colonizers had a cell-density growth requirement when inoculated as monocultures. All strains successfully grew in mucin-serum even when inoculated at a cell-density as low as 10^3^ cells mL^−1^ (Figure 2f), which suggests these species are able to grow from small populations without requiring an autoinducing factor.

### *Veillonella parvula* is the key species that supports growth of low-cell-density *P. gingivalis*

Evaluation of the individual ability of each of the five early colonizing species to support growth of a low-cell-density inoculum of *P. gingivalis*, showed that only *V. parvula* enabled the latter to grow (Figure 3a). Furthermore, when all microorganisms were inoculated as a community, elimination of *V. parvula* from the inoculum abrogated growth of *P. gingivalis* (Figure 3b), confirming *V. parvula* as the key species.

**Figure 3.**
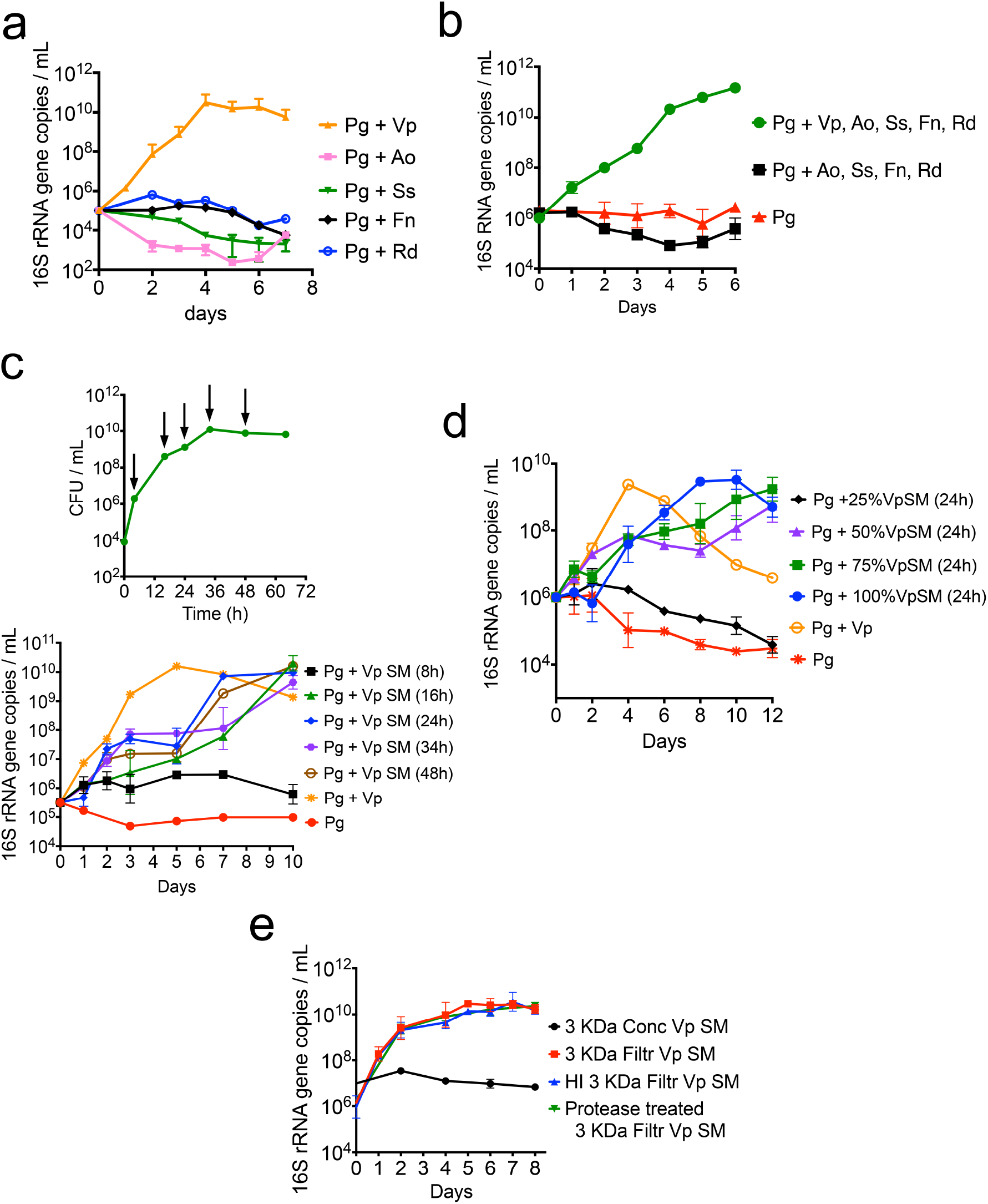
*V. parvula* (Vp) is the key species that through a diffusible factor supports growth of low-cell-density *P. gingivalis* (Pg). **a.** Growth of Pg when co-inoculated (at 10^5^ cells mL^−1^) in mucin-serum with either Vp, *Actinomyces oris* (Ao), *Streptococcus sanguinis* (Ss), *Fusobacterium nucleatum* (Fn) or *Rothia dentocariosa* (Rd). Graph shows Pg growth as determined via qPCR. **b.** Presence of Vp is essential for the growth of a low cell-density inoculum of Pg. Graph shows Pg growth, as determined via qPCR, when inoculated together with all initial colonizers, in the absence of Vp, or as a monoculture. **c.** Evaluation of the effect of Vp spent medium (SM) on Pg growth. SM was collected from a Vp batch culture in mucin-serum at the times indicated by the arrows (green curve, top panel), filtered and used to evaluate growth of Pg (10^5^ cells mL^−1^). **d.** Evaluation of the effect of different concentrations of Vp SM (collected at 24 h) on growth of a low-cell-density Pg inoculum showing dose-dependent stimulation of growth by Vp SM. **e.** Soluble factor in Vp SM capable of supporting growth of low-cell-density Pg is smaller than 3 kDa, is heat-stable and is protease resistant. SM from Vp grown for 24 hours in mucin-serum was filtered through 3 kDa membranes and either heat-inactivated (HI) or treated with proteases, followed by lyophilization and reconstitution (10x) in dIH20. Reconstituted fractions (Conc = > 3kDa and Filtr = <3kDa) were added to fresh mucin-serum medium (1:3, vol:vol) to evaluate growth of low-cell-density Pg (10^5^ cells mL^−1^).

The positive effect of *V. parvula* on *P. gingivalis* was strain-independent as three different *P. gingivalis* strains (W83, 381 and ATCC 33277) grew when co-inoculated with any of three different strains of *V. parvula* (PK1910, PK1941 and ATCC 10790) but failed to grow as monocultures (Supplemental Figure 2).

### *V. parvula* supports growth of a low-cell-density inoculum of *P. gingivalis* through a soluble signal

Spent medium from *V. parvula*, collected after different lengths of time in culture, was evaluated for its ability to stimulate growth of a low-cell-density inoculum of *P. gingivalis*. As shown in Figure 3c, spent medium from early cultures (8 hours) did not support growth of *P. gingivalis* but that obtained from cultures older than 16 h supported growth, although the growth rate in spent medium was slower than when *V. parvula* was present. The effect of spent medium from *V. parvula* was clearly dose-dependent since addition of 25% spent medium to fresh medium did not allow growth of *P. gingivalis*, while 50% or higher concentrations supported growth (Figure 3d). Cell-to-cell contact was not essential for the interaction since separation of *V. parvula* and *P. gingivalis* by a 0.22 μm filter still allowed the latter species to grow (Supplemental Figure 3a). We also noticed that if only spent media from *V. parvula*, but not cells, were allowed to interact with *P. gingivalis* (as in Figures 3c, 3d and Supplemental Figure 3a), biphasic growth tended to occur with a slight decrease in growth rate as *P. gingivalis* approached the threshold concentration needed to sustain its own growth. This biphasic growth was not observed with a larger inoculum (Supplemental Figure 3b, left panel). Spent medium of *V. parvula* was also seen to have no effect on inocula capable of independent growth (Supplemental Figure 3b, right panel).

The spent medium of *V. parvula* showed similar characteristics to the auto-stimulatory spent medium of *P. gingivalis*. That is, only the <1kDa filtrate fraction enabled growth of *P. gingivalis* (Supplemental Figure 3c), and the activity of the filtrate was heat-stable and protease resistant (Figure 3e).

In summary, these results suggest *V. parvula* produces a diffusible small molecule that needs to accumulate to a threshold concentration to stimulate growth of low-cell-density *P. gingivalis*. Although cell-free spent medium supported growth, presence of *V. parvula*, but not necessarily cell-to-cell contact, was beneficial to the interaction.

### *V. parvula* allows *P. gingivalis* to maintain a high biomass in an open flow chemostat system

To evaluate if the interaction between *V. parvula* and *P. gingivalis* was relevant in an open flow setting, which more closely resembles the conditions in natural oral communities, we used a continuous culture system. For these experiments, early colonizers and *P. gingivalis* were inoculated into a chemostat, allowing microorganisms to briefly grow in batch before turning on the flow of growth medium. Since our objective was to evaluate whether *V. parvula* helped *P. gingivalis* maintain its biomass under open flow, we used a high-density inoculum during the culture establishment. *P. gingivalis* was able to grow in continuous culture in the absence of *V. parvula* (Figure 4a), but its steady-state biomass was significantly higher when *V. parvula* was part of the initial inoculum (Figure 4b) or introduced after steady-state (Figure 4c). *V. parvula* not only allowed *P. gingivalis* to reach higher cell numbers in this open-flow system, but it also reduced daily biomass fluctuations of *P. gingivalis* (Figure 4d). These results suggest that *V. parvula* is also beneficial to high-cell-density *P. gingivalis* enabling it to maintain a higher biomass under open-flow conditions.

**Figure 4.**
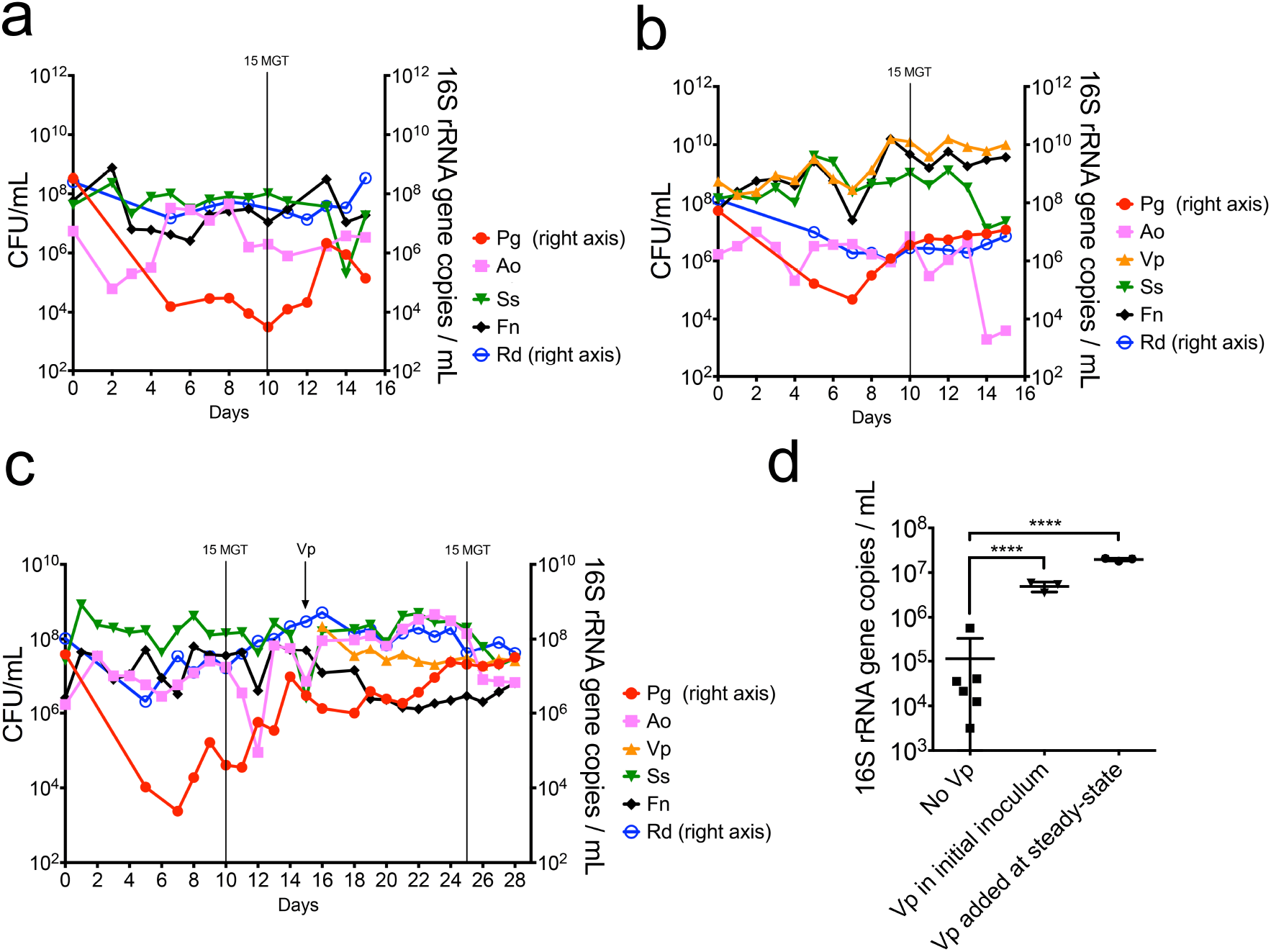
*V. parvula* (Vp) helps *P. gingivalis* (Pg) maintain a high biomass when growing as part of a polymicrobial community under open-flow continuous-culture conditions. **a**. Pg was co-inoculated in a chemostat in mucin-serum with *Actinomyces oris* (Ao), *Streptococcus sanguinis* (Ss), *Fusobacterium nucleatum* (Fn) and *Rothia dentocariosa* (Rd). **b.** Vp was added to the initial inoculum together with Pg, Ao, Ss, Fn and Rd. **c**. Pg was initially co-inoculated with Ao, Ss, Fn and Rd (in the absence of Vp) and the culture was allowed to achieve steady-state, after which Vp was added. **d.** Direct comparison of Pg biomass at steady-state (including 3 time points after 15 mean generation times, MGT) in the absence and presence of Vp. In all experiments, a high density (10^8^ CFU/mL) inoculum was employed for all species. Cell numbers of Vp, Ao, Ss and Fn were determined by plating on selective media. Biomass of Pg and Rd was determined via qPCR. **** represents P < 0.0001 after t-tests.

### Low-cell-density *P. gingivalis* is unable to colonize the oral cavity of mice unless aided by *V. parvula*

The ligature-induced periodontitis (LIP) mouse model was used to evaluate whether the requirement for a high-cell-density inoculum was relevant in an in vivo oral environment and to explore if under these conditions *V. parvula* had a positive effect on colonization by low cell-density *P. gingivalis*. In this model, ligatures that promote accumulation of bacteria are tied around molar teeth leading to bacterial dysbiosis and an inflammatory process that induces bone loss within 5 days. A small volume (50 μL) of a *P. gingivalis* suspension was inoculated on the ligatures, at the time of placement, at a low (10^5^ cells mL^−1^) or high (10^8^ cells mL^−1^) cell-density. Five days post-inoculation, *P. gingivalis* was only detected when inoculated at high cell-density (Figure 5a). However, the inability of low-cell-density *P. gingivalis* to colonize was reversed when *V. parvula* was co-introduced in the inoculum, enabling low-cell-density *P. gingivalis* to reach similar colonization levels to those achieved by *P. gingivalis* when introduced alone at high cell-density (Figure 5a). Furthermore, *V. parvula* significantly enhanced the ability of high cell-density *P. gingivalis* to colonize, in comparison to high-cell-density *P. gingivalis* when introduced alone (Figure 5a).

**Figure 5.**
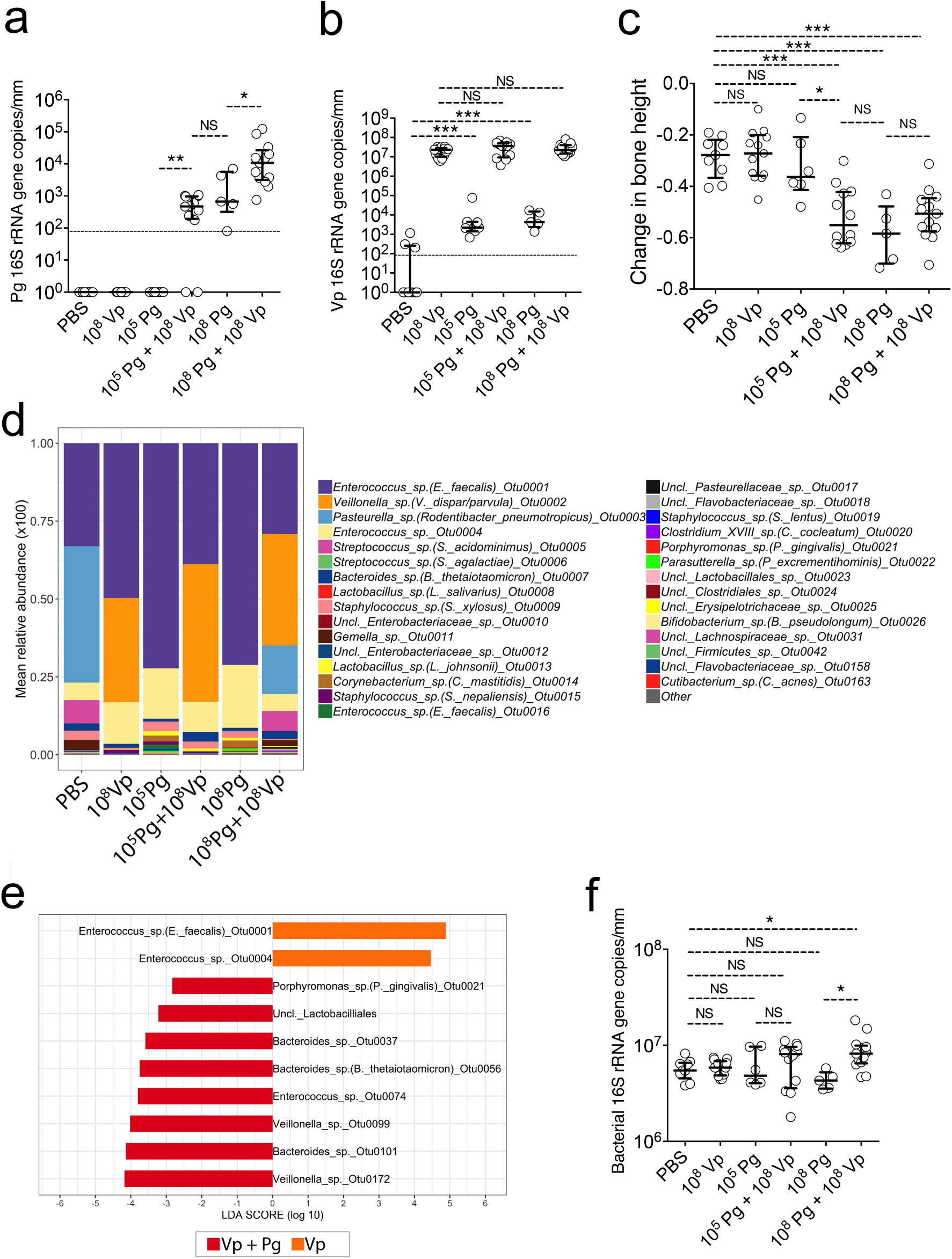
*V. parvula* (Vp) allows a low-cell-density inoculum of *P. gingivalis* (Pg) to colonize and augment bone loss in a ligature-induced periodontitis murine model. **a.** Pg levels measured via qPCR on ligatures retrieved 5 days post-inoculation. Horizontal line shows limit of detection of the assay. **b.** Vp levels of retrieved 5-day ligatures as evaluated via qPCR. Horizontal line shows limit of detection of the assay. **c.** Alveolar bone levels after 5 days of ligature placement and inoculation. **d.** Microbiome composition of retrieved 5-day ligatures as evaluated via 16S rRNA gene sequencing. **e.** LEfSe evaluation of operational taxonomic units (OTUs) with different relative abundance when Vp was inoculated alone in contrast to Vp co-inoculated with Pg. **f.** Total bacterial load of retrieved 5-day ligatures as evaluated via qPCR and universal primers. *** indicates a p value <0.001, ** indicates a p<0.01 and * a p<0.05 (Mann-Whitney Rank tests). NS= not statistically significant.

Figure 5b shows the levels of *V. parvula* in the different groups. *V. parvula* was a very efficient colonizer reaching similarly high numbers when introduced alone or with *P. gingivalis*. It was also observed that in the groups in which *V. parvula* was not exogenously introduced, there was a low number of indigenous *V. parvula* present. This is an expected finding as *V. parvula* has been shown to be a minor component of the oral microbiome of certain strains of mice (*22*). These basal low levels of indigenous *V. parvula*, however, were insufficient to facilitate colonization of *P. gingivalis*, in agreement with our in vitro culture results, which indicated that that a minimum threshold biomass of *V. parvula* is needed to stimulate growth of *P. gingivalis*.

### Colonization of *P. gingivalis* promotes periodontal pathology (bone loss)

We next assessed the consequences of *V. parvula* and *P. gingivalis* oral colonization by measuring periodontal bone loss in the mice subjected to LIP and locally inoculated, or not, with these bacteria separately or in combination. The colonization of *V. parvula* alone did not increase bone loss in comparison to the PBS negative control group (Figure 5c), confirming that *V. parvula* is normally a symbiotic commensal. In contrast, in mice in which colonization of *P. gingivalis* occurred, either because it was introduced at high-cell-density or at low-cell-density aided by *V. parvula*, there was significantly greater bone loss compared to the PBS control group, in which bone loss is driven solely by dysbiotic indigenous bacteria. Introduction of *P. gingivalis* in the oral cavity of healthy mice (not subjected to LIP) has been shown to cause microbiome dysbiosis and an increase in the total bacterial load (*13*). To examine if *P. gingivalis* induced greater dysbiosis than that already occurring due to ligature placement, the microbiome composition was determined (Figure 5d). In agreement with the qPCR observations, the 16S rRNA gene data confirmed *P. gingivalis* to be as a minor component of the community, while *V. parvula* occupied about a third of the total biomass when exogenously introduced (Figure 5d). The overall community composition, however, was not dramatically modified by the introduction of *P. gingivalis* but some changes in low-abundance species occurred. When comparing species differentially enriched in mice inoculated with *V. parvula* and *P. gingivalis* versus those inoculated with *V. parvula* alone, it can be seen that in the presence of *P. gingivalis*, a few low-abundance species, including other members of the *Bacteroidetes* phylum, became enriched, while certain *Enterococcus* spp. were depleted (Figure 5e). In contrast to these compositional changes, *P. gingivalis* colonization was not associated with a higher bacterial load (Figure 5f).

In summary, the symbiotic commensal *V. parvula* enabled a pathogenic species, *P. gingivalis*, to colonize the mouse oral cavity and augment bone loss. In this model, colonization of *P. gingivalis* increased the abundance of some minor constituents, such as other *Bacteroidetes*, although it did not affect the total bacterial load or dramatically altered the microbiome community composition.

## Discussion

In this study, we show that the growth of a pathogen that is implicated in the etiology of the oral disease periodontitis depends on cell-density, which determines the concentration of an endogenous soluble small molecule that is essential for growth. Such an inability to grow from a low cell-density population in vitro was also observed in vivo, as *P. gingivalis* was unable to colonize the oral cavity of mice when introduced at low cell-density. The requirement for this autoinducing soluble factor may restrict the colonization of *P. gingivalis* in the human oral cavity. This is consistent with the low detection of *P. gingivalis* in periodontal health, as shown by this study, and its unstable colonization in young individuals (*14*). Periodontitis, however, is a prevalent condition with severe disease affecting about 10% of the global population (*23*). As shown in our re-analysis of publicly available subgingival microbiome datasets, about 70% of subjects with periodontitis had detectable *P. gingivalis*. Among those individuals, about 60% had *P. gingivalis* at greater than 1% relative abundance. This sets a scenario in which transmission of this pathogen from humans with severe periodontitis to other unaffected hosts is likely to occur frequently, but colonization of recipients is limited by the inability of *P. gingivalis* to grow from low cell-density inocula.

Under specific circumstances, however, such as in the presence of undisrupted dental biofilm accumulation, a specific cross-species interaction with a ubiquitous early-colonizing species, *V. parvula*, may allow establishment of *P. gingivalis* in the human oral cavity. *V. parvula* is one of the earliest colonizers of tooth surfaces and becomes a dominant community component as dental biofilms mature (*4, 15*). *V. parvula* is also a core subgingival species, that is, a species that is present at equal relative abundance in both health-associated and dysbiotic microbiome communities (*3*). However, since the total community biomass is higher in disease, the total load of *V. parvula* increases in the dysbiotic state. Such increase in biomass of this commensal species during plaque maturation seems to be a key component of the interaction with *P. gingivalis*, since both in vitro and mouse experiments showed that th mere presence of *V. parvula* was insufficient to enable growth of low-cell-density *P. gingivalis*, but instead, high cell numbers of *V. parvula* were needed. Therefore, the accumulation of *V. parvula* in dental biofilms, such as those associated with the gingivitis state, may allow establishment of *P. gingivalis* since the soluble factor provided by *V. parvula* would only then reach a threshold concentration for growth of the pathogen. Gingivitis also leads to sporadic bleeding upon tissue stimulation (eg. during tooth brushing) and, as we show here, blood enables growth of *P. gingivalis* from even small inocula. The inflammatory exudate present during gingivitis also creates the necessary nutritional environment propitious for the growth of the peptide-dependent and proteinase-rich *P. gingivalis* (*16*). Accordingly, here we observed a change in detection of *P. gingivalis* from about 1% of subjects in health to about 25% in gingivitis (Fig. 2a). Moreover, as community maturation progresses towards the periodontitis state, *V. parvula* would still be beneficial to *P. gingivalis* helping it maintain a high biomass as indicated by our chemostat and LIP experiments. A model depicting the interaction between *V. parvula* and *P. gingivalis*, as mediated by the accumulation of soluble small diffusible molecules during dental biofilm maturation is shown in Figure 6.

**Figure 6.**
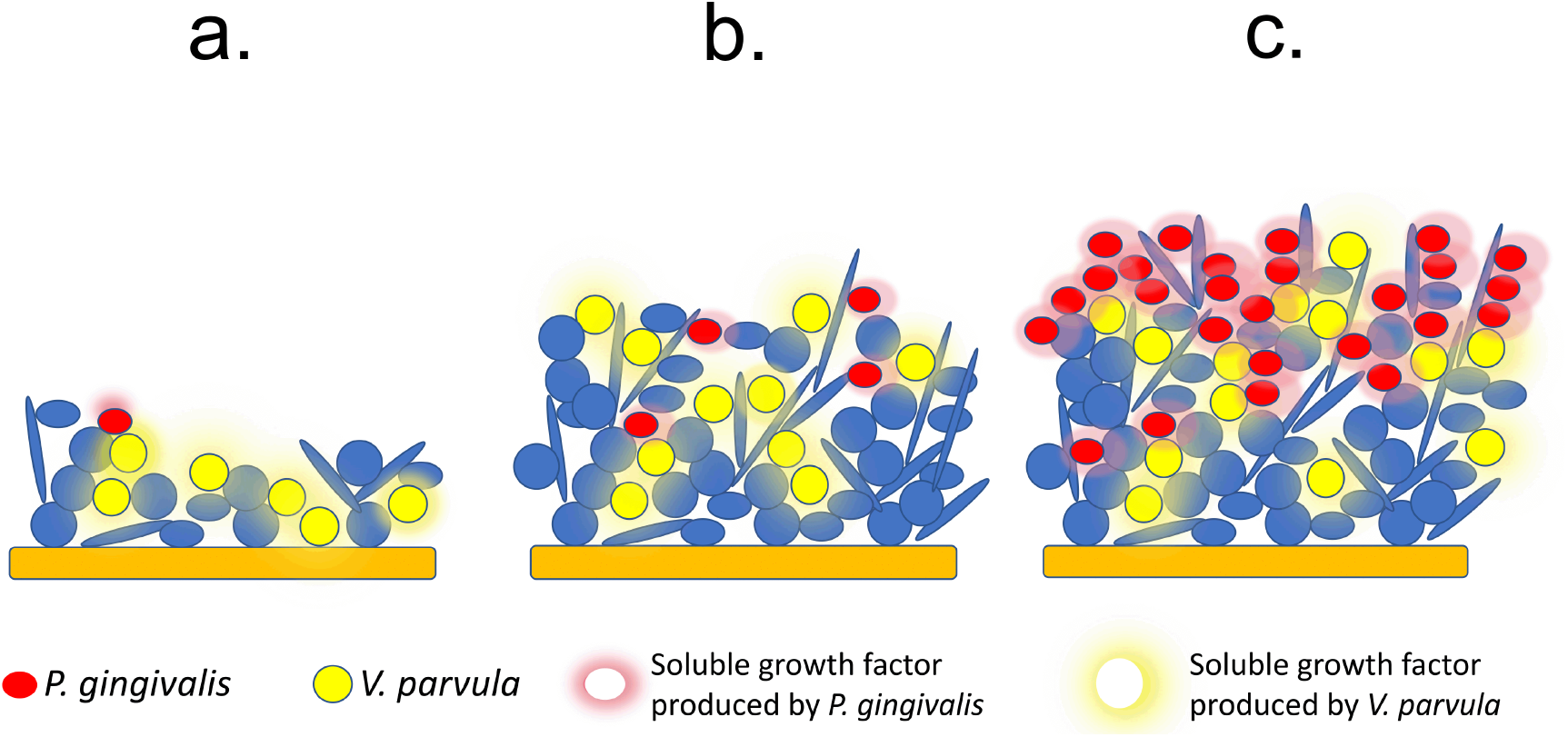
Model depicting *V. parvula* (Vp)*-P. gingivalis* (Pg) interaction during dental biofilm community development. **a.** During early stages of biofilm formation on tooth surfaces, Pg is not able to establish since it cannot grow from a low-cell-density population. Vp does not depend on cell-density so it can grow and become established during early stages of biofilm maturation. **b.** If dental communities are left undisturbed, as is the case in gingivitis, Vp increases in biomass, producing a low-mass soluble factor that accumulates to a threshold concentration capable of supporting growth of Pg. **c.** Once Pg becomes established at high-cell-density, such as in a dysbiotic biofilm associated with periodontitis, its growth is supported by its own soluble low-mass growth factor. Vp, which is also an abundant species in mature plaque (core species) contributes to stabilizing Pg biomass in the dysbiotic periodontitis-associated community.

Although *P. gingivalis* and *V. parvula* are able to coaggregate together, physical contact was not needed for growth stimulation to occur in a closed culture system. The presence of *V. parvula* in co-cultures, however, was beneficial and allowed for faster growth of *P. gingivalis*, compared to growth in spent media of the former species. It is then possible that in natural oral communities, which are an open-flow environment, close cell-to-cell distance benefits the inter-species interaction described here.

Characterization of spent media from *P. gingivalis* and *V. parvula* showed that the growth-inducing activity originates from a non-proteinaceous, heat-stable, small molecule, and is, therefore, similar in nature to other quorum-sensing mediators (*1*). Although much more work is required to identify the molecule(s) responsible for inducing growth, our data suggest that the nature of the interaction between *P. gingivalis* and *V. parvula* is unidirectional (commensalism) as no obvious benefit was seen for the latter species. In accordance with this, the molecule mediating the interaction could be considered in an evolutionary context as a ‘cue’, and not a ‘signal’ (*1*), as *V. parvula* does not seem to directly benefit from interacting with *P. gingivalis*. In contrast, *P. gingivalis* exploits several aspects of the metabolism of *V. parvula*. Apart from providing a growth-initiating factor, *V. parvula* has been shown to produce heme which is the preferred iron source for *P. gingivalis* (*24*). *Veillonella* is also able to detoxify hydrogen peroxide produced by other early colonizing species via its catalase activity, facilitating growth of less oxygen-tolerant anaerobes (*8*). Therefore, *Veillonella* interacts with *P. gingivalis* via distinct mechanisms that collectively support the establishment of the latter in the gingival crevice. In this context, therefore, *V. parvula* behaves as an ‘accessory pathogen’, *i.e*., an organism that, while commensal in a particular microenvironment, can also support or augment the colonization and/or virulence of pathogenic microorganisms (*25*).

Different types of inter-species interactions occur in polymicrobial communities. While two-species interactions are the simplest type, in natural environments interactions can occur among several species, generating indirect and emergent effects. Despite this potential complexity, here we demonstrate that the dual species interaction between *V. parvula* and *P. gingivalis* was relevant within the context of a community; and although we do not exclude the possibility that higher order interactions also took place, the pairwise interaction remained valid in a polymicrobial context. In one of the community models tested *in vitro* (the 6-species community), it was shown that member species other than *V. parvula* did not directly interact with *P. gingivalis,* and therefore, *V. parvula* was the only relevant partner. However, in the mouse oral cavity, the interactions of local commensals and *P. gingivalis* were uncertain and yet, low-cell-density *P. gingivalis* was unable to grow when inoculated alone, whereas *V. parvula* was able to support its colonization. These findings highlight the importance of the specific interaction discovered and show that it is possible to identify key pairwise inter-interactions within complex communities. It would be relevant to investigate if other human dental plaque species, different to the early colonizers tested here, are able to support growth of *P. gingivalis*, since our work suggests that limiting the biomass of potential accessory pathogens, such as *Veillonella*, may preclude *P. gingivalis* from colonizing the oral cavity. Excluding *P. gingivalis* could in turn block the growth of other species within pathogenic communities whose growth depends on the immune dysregulation induced by *P. gingivalis* (*13*).

Experiments using the mouse LIP model show that *P. gingivalis* colonization of an already dysbiotic microbiome augmented bone loss, confirming its pathogenic capacity, in contrast to *V. parvula*, which did not have any effect in that regard. In another mouse model of *P. gingivalis*-induced bone loss, the microorganism is inoculated via swabs into orally healthy mice (*26*). In that oral inoculation model, repeated introduction of *P. gingivalis* leads to its establishment as a low-abundance member of the microbiome that induces bone loss by manipulating the host inflammatory response and undermining the effectiveness of immune bacterial clearance (*12, 13*). Thus, in the oral inoculation model, *P. gingivalis* acts as a keystone pathogen leading to an increase in the whole commensal community biomass and to qualitative changes in community composition, which is thought as the cause of bone loss (*13*). There are some similarities between that oral inoculation model (*26*) and our current work. In the LIP model used in our study, *P. gingivalis* also became a minor constituent (about 0.1%) of the total community biomass, augmenting bone loss beyond that already produced by ligature placement alone. However, in contrast to the oral inoculation model, here *P. gingivalis* did not cause an increase in bacterial load or dramatic compositional changes to the microbiome beyond those already present. In other words, whereas *P. gingivalis* induces *de novo* alterations to the community structure in the oral inoculation model (*13*), it does not appear to profoundly enhance LIP-induced dysbiosis. However, *P. gingivalis* colonization caused an enrichment of some minor microbiome components, including other *Bacteroidetes* species, the significance of which in any pathogenic process is currently uncertain. The two models therefore agree in that *P. gingivalis* can promote bone loss even as a low-abundance community member, but the mechanisms by which *P. gingivalis* augments bone loss in the two models may be different.

Although the tooth-associated subgingival biofilm is the predominant habitat of *P. gingivalis*, this pathogen and virulence factors thereof have been localized in remote tissues in association with comorbid conditions (*6*). For instance, in Alzheimer’s disease, *P. gingivalis* was shown to ectopically infect the brain of humans and mice and to correlate with or cause neuronal pathology, in humans and mice, respectively (*27*). Therefore, approaches to control the growth of *P. gingivalis* in its predominant habitat (which serves as a reservoir for its systemic dissemination) may also help reduce the risk of periodontal comorbidities in which *P. gingivalis* is implicated. Our work provides a novel target to control the growth of *P. gingivalis* (and hence its dissemination); although indirect, successful targeting of the accessory pathogen *V. parvula* should prevent the ability of *P. gingivalis* to expand within the oral microbial community to levels at which it can become pathogenic.

Altogether, our work demonstrates that a requirement for a cell-density dependent signal limits growth and colonization of the human oral pathogen *P. gingivalis*. This inability to establish from a small inoculum is overcome by forming a specific partnership with a ubiquitous commensal species of human dental biofilms, which after increasing in biomass is able to provide the growth-initiating signal at the concentration required by *P. gingivalis*. These results shed light into some of the mechanisms behind dental biofilm microbial successions and highlight the role of cell-density-mediated interactions between early- and late-colonizers ultimately leading to pathogen colonization and virulence.

## Methods

### Strains and culture conditions

*Porphyromonas gingivalis* strains 381, W83 and ATCC 33277 were maintained short-term on agar containing brain heart infusion (BHI), 0.04% L-cysteine ·HCl, 5 ug mL^−1^ hemin, 5 μg mL^−1^ menadione and 5% defibrinated sheep’s blood. Starter cultures were grown in BHI, 0.04% L-cysteine ·HCl, 5 μg mL^−1^ hemin and 5 μg mL^−1^ menadione (BHI-H-M). *Streptococcus sanguinis* SK36 and *Actinomyces oris* T14V were maintained on BHI agar and grown in liquid BHI. *Fusobacterium nucleatum* subsp. *polymorphum* ATCC 10953 was maintained on agar containing BHI, 0.04% L-cysteine ·HCl and 5% defibrinated sheep’s blood and starter cultures were grown in BHI and 0.04% L-cysteine ·HCl. *Veillonella parvula* strains PK1910, PK1941 and ATCC 10790 were maintained on agar containing BHI, 0.04% L-cysteine ·HCl and 1.3% lactic acid and starter cultures were grown in a similar liquid medium. Cultures of the previous microorganisms were incubated in an anaerobic atmosphere consisting of 5% H_2_, 5% CO_2_, and 90% N_2_. *Rothia dentocariosa* ATCC 17931 was grown on BHI agar or in liquid BHI aerobically. The strain *P. gingivalis* Δ*luxS*::*ermF* (*21*), kindly donated by Dr. Richard J. Lamont, University of Louisville, was maintained in the presence of erythromycin at 15 μg mL^−1^.

### Evaluation of growth from inocula of varying cell-densities

To evaluate the ability of inocula of different size to grow, microorganisms were inoculated into liquid cultures at cell densities ranging from 10^3^ to 10^8^ cells per mL followed by anaerobic incubation at 37°C. Inocula of different biomass were obtained by diluting a starter culture previously grown to mid logarithmic phase and normalized to an optical density (600 nm) of 0.4, for which the number of cells was determined according to microscopic counts on a Petroff Hausser chamber. Most experiments were conducted in mucin-serum medium, which contained 2.5 mg mL^−1^ hog gastric mucin (Sigma), 2.5 mg mL^−1^ KCl, 2.0 mg mL^−1^ proteose peptone, 1.0 mg mL^−1^ yeast extract, 1.0 mg mL^−1^ trypticase peptone, 1.0 μg mL^−1^ cysteine ·HCl, 5 μg mL^−1^ hemin and 10% (vol:vol) heat-inactivated human AB serum (Sigma). Cultures were sampled daily, inside the anaerobic chamber. Growth was monitored after serial dilutions and plating on appropriate media or evaluated via qPCR.

### Evaluation of the effect of cell-free spent medium on growth of *P. gingivalis*

Spent medium was obtained from *P. gingivalis* cultures inoculated with 10^7^ cells mL^−1^ and grown until late exponential phase (48 hours in BHI-H-M and 72 hours in mucin-serum), followed by centrifugation for 15 min at 5,000 × g. Supernatants were filter-sterilized twice through 0.22 μm filter units. The filtered spent medium was checked for contamination by plating a small volume on blood agar followed by aerobic and anaerobic incubation. Spent medium was stored at 4°C for up to 48 h before using it to evaluate the growth of *P. gingivalis*. The effect of spent media was tested by combining it in different proportions (25 to 100%) with fresh medium, followed by inoculation of *P. gingivalis* at low-cell-density (10^5^ cells mL^−1^). To evaluate the effect of spent media on solid growth on agar, *P. gingivalis* was serially-diluted in either PBS or spent medium, and plated at different densities onto BHI-H-M or BHI-H-M blood agar. Colonies were counted after anaerobic growth for 8 days.

*V. parvula* spent medium was obtained at different time points of growth in mucin-serum (with most experiments ultimately conducted with spent medium from 24-hour cultures). Spent medium was processed in a similar manner to that described for *P. gingivalis*.

### Fractionation, heat inactivation and protease treatment of spent medium

Spent medium was subjected to fractionation based on size of molecules by using spin filter units with different molecular weight cut-offs (MWCO) (Amicon® Ultra or Microsep ™ Advance centrifugal devices). Samples were loaded and centrifuged at 5,000 × g for 90 min and concentrates and filtrates were freeze-dried followed by reconstitution in dIH_2_O giving 10X concentrated suspensions. These concentrated fractions were tested for their ability to induce growth of a low-cell-density inoculum of *P. gingivalis* by adding them to fresh mucin-serum (25% -75% vol:vol). Concentrated low molecular weight filtrates were heat-inactivated by boiling for 10 min and then cooled before using them to evaluate growth of *P. gingivalis*. Filtrate fractions were protease-treated by incubation, for 1 h, with 2 U mL^−1^ of reconstituted proteinase K-agarose dry powder (Sigma). Protease-treated fractions were then centrifuged to remove proteinase K-agarose and the supernatant recovered and used to evaluate growth of *P. gingivalis*.

### Evaluation of the effect of quorum-sensing-related compounds on growth of *P. gingivalis*

A group of commercially available compounds potentially involved in cellular growth induction, including D-pantothenic acid (D-PA), D-panthenol, β-alanine, tyrosol, as well as the polyamines spermidine, spermine, cadaverine and putrescine were tested for their ability to induce growth of a low cell-density inoculum of *P. gingivalis* (10^5^ cells mL^−1^). *P. gingivalis* was inoculated in mucin-serum medium supplemented with each compound at the concentrations listed in Supplementary Table 1, and growth was monitored for up to 10 days. A high-cell-density inoculum (10^8^ cells mL^−1^) of *P. gingivalis* and a co-culture of 10^5^ cells mL^−1^ *P. gingivalis* and 10^5^ cells mL^−1^ *V. parvula,* placed in unsupplemented mucin-serum were included as positive controls.

### Evaluation of the effect of cell-to-cell contact with *V. parvula* on growth of *P. gingivalis*

Mucin-serum aliquots inoculated with *P. gingivalis* and/or *V. parvula* at 10^5^ cells mL^−1^ were placed into 50 mL conical tubes separated by a 0.22 μm membrane (Steriflip-GP, Millipore). Three conditions were tested: (i) *P. gingivalis* monoculture inoculated in one chamber and *V. parvula* monoculture in the contiguous one, (ii) *P. gingivalis* + *V. parvula* inoculated in both chambers, and (iii) *P. gingivalis* inoculated in both chambers. Cultures were sampled and *P. gingivalis* growth was evaluated via qPCR.

### Evaluation of the effect of early colonizers on the growth of *P. gingivalis* in batch and continuous culture

*P. gingivalis* was inoculated in batch as a monoculture or in the presence of early colonizers in mucin-serum. Inoculum size was 10^5^ cells mL^−1^ for all species. Cultures were sampled daily inside the anaerobic chamber. Growth of all species was monitored after serial dilution and plating on selective agar media or evaluated via qPCR.

Continuous culture experiments were performed in a Bioflow®/CelliGen® 115 Bioreactor (New Brunswick) starting from standardized frozen inocula stored in medium specific for each microorganism and 10% glycerol. At inoculation, cryovials containing standarized stocks were rapidly allowed to thaw, followed by pooling of different strains and inoculation into 500 mL of mucin-serum. Inoculation density was 10^8^ cells mL^−1^ for each strain. After 24 hours of batch growth in the bioreactor vessel, the pump was turned on and fresh medium allowed to flow for 48 hours at a dilution rate of *D*=0.0462 h ^−1^ (doubling time *t*d=15 h). The flow was then stopped and a new inoculation was performed, followed by batch growth for 24 hours, after which continuous culture was resumed. This time point was considered day 0. The gas phase was maintained anaerobically by sparging 5% CO_2_ in N_2_; temperature and pH were controlled automatically at 37°C and 7.15 ± 0.15, respectively. Cultures were considered to have reached steady state after 15 mean generation times (MGT), and evidence of sustained stability as evaluated via dry weights, *E* h and viable counts. Three different types of experiments were conducted to evaluate the effect of *V. parvula* on the biomass of *P. gingivalis*. In one experiment, *A. oris*, *S. sanguinis*, *F. nucleatum* and *R. dentocariosa* were inoculated together with *P. gingivalis*. In a second set of experiments, *V. parvula* was included in the initial inoculum in addition to the strains already mentioned. In a third set of experiments, *V. parvula* was initially excluded but introduced later once the culture had achieved steady-state, after which the culture was monitored until a second steady-state was reached.

### Cultivation and molecular methods for quantification of microorganisms from batch and continuous cultures

Growth of *S. sanguinis*, *A. oris*, *V. parvula* and *F. nucleatum* were quantified by plating on appropriate selective media. Culture samples were vortexed, followed by a 10s sonication at 15% amplitude in a Branson sonicator model 4C15, to disperse co-aggregated microbial cells without affecting viability. After disagreggation, appropriate dilutions in sterile phosphate buffered saline (PBS) were obtained and subsequently plated. BHI supplemented with 5% defibrinated sheep’s blood, 0.04% L-cysteine ·HCl and 0.0025 g L^−1^ vancomycin hydrochloride was used to quantify *V. parvula* and *F. nucleatum* (anaerobically), differentiating them by colony morphology. Actino-selective agar consisting of trypticase soy agar supplemented with 0.5% glucose, 0.0013% cadmium sulfate, 0.008% sodium fluoride, 0.00012% neutral acriflavine, and 0.000025% basic fuschin was used to quantify *A. oris* (anaerobically). Mitis-Salivarius agar was used to quantify *S. sanguinis* (aerobically). *P. gingivalis*, and *R. dentocariosa*, were quantified by qPCR. The reason for using a molecular technique to quantify *R. dentocariosa* is that no suitable selective medium was found for its identification. qPCR was also more reliable to quantify *P. gingivalis*, especially when the microorganism was present in low-abundance as part of a multi-species community. For these assays, DNA extraction from cultures was performed as previously described (*3*)*. P. gingivalis* and *R. dentocariosa* were quantified using primers targeting the 16S rRNA gene, and amplicons detected via SYBR green chemistry or a Taqman probe, respectively (*28, 29*). Standard curves using the respective genomic DNA were used to calculate number of 16S rRNA gene copies present in samples.

### Reanalysis of 16S rRNA gene amplicon libraries of subgingival samples obtained from subjects with different periodontal conditions

An evaluation of the prevalence and abundance of *P. gingivalis* and early colonizers in subjects presenting with periodontal health, gingivitis and periodontitis was performed. Integration and re-analysis of datasets from different published studies was required since no simultaneous analysis of the microbiome of these three conditions, allowing direct comparison and applying current clinical definitions, has been reported. Studies that used 16S rRNA gene amplicon sequencing of the V1-V3 hypervariable regions to characterize the subgingival microbiome in health, gingivitis or periodontitis; and with downloadable publicly available sequence datasets were included (*3–5, 30–36*). Sample selection from these studies was based on their compatibility with current definitions of periodontal health, gingivitis and periodontitis. Studies included in the periodontal health group were required to exclude subjects with >10% bleeding on probing and pocketing >3 mm. Studies included in the gingivitis group enrolled subjects with naturally-occuring gingivitis defined by >10% bleeding on probing but no pocket ≥5 mm, or periodontally-healthy subjects who underwent an experimentally-induced gingivitis protocol. Subjects with periodontitis were included based on the minimum case definition for the disease, which is interdental clinical attachment loss (CAL) detectable at ≥2 non-adjacent teeth, or buccal CAL ≥3 mm with pocketing >3 mm detectable at ≥2 teeth. Only those samples from subjects who were non-smokers, non-diabetic and that did not have chronic kidney disease were included in the analysis.

Downloaded sequences were processed in mothur, using standard pipelines (*37*). Sequences were classified to species level by using the classify.seqs command and the Human Oral Microbiome database (HOMD) V14.5 as reference. After processing, sequence libraries were randomly subsampled at a threshold of 3,500 reads to contain the same number of reads followed by generation of relative abundance tables.

### Effect of *V. parvula* on the in vivo colonization and virulence of *P. gingivalis*

All animal experiments were reviewed and approved by the Institutional Animal Care and Use Committee (IACUC) of the University of Pennsylvania and were performed in compliance with institutional, state, and federal policies. A previously described ligature-induced periodontitis (LIP) mouse model was used (*38*), modified to include inoculation of exogenous microorganisms. Briefly, ligatures were tied around molar teeth of 8 week-old C57BL/6 mice and 50 μL of a suspension, in phosphate buffered saline (PBS), of 10^5^ cells mL^−1^ or 10^8^ cells mL^−1^ of *P. gingivalis*, *V. parvula* or a combination of both was placed directly on the ligatures. Only one inoculation, at the time of ligature placement, was performed. Ligatures were removed 5 days post-placement and alveolar bone levels were evaluated as previously described (*38*).

DNA was extracted from ligatures using a previously described protocol (*3*). Total bacterial load was determined by qPCR using universal primers and a TaqMan probe (*3*). *P. gingivalis*, and *V. parvula* load was determined using specific primers targeting the 16S rRNA gene and detected via SYBR green chemistry or a TaqMan probe, respectively (*29, 39*). Standard curves were used to calculate number of 16S rRNA gene copies in each condition. Data were expressed as 16S rRNA copy number normalized by ligature length. Microbiome communities in ligatures were characterized by sequencing of the 16S rRNA V1-V2 region using primers 8F 5’- AGAGTTTGATCMTGGCTCAG-3’ and 361R 5’- CYIACTGCTGCCTCCCGTAG-3’ which included the adapter for MiSeq sequencing (Illumina) and single end barcodes (*4*). Amplicon libraries were pooled and sequenced using the MiSeq Reagent kit v3 (Illumina). 16S rRNA gene sequences were processed in mothur using standard pipelines. Reads were clustered at 97% similarity into Operational Taxonomic Units (OTUs). Individual reads were classified by comparison to the RDP version 16 database, as implemented in mothur, with a cutoff=80. OTUs were classified up to genus level when possible, according to the consensus taxonomy using the default cutoff (51%). To enhance the taxonomical resolution of each OTU, the representative sequence was compared using BLAST to the NCBI 16S rRNA gene sequence database and the best match (with at least 97% similarity and coverage) is indicated in parenthesis. Relative abundance graphs were generated using the packages ‘ggplot2’ and ‘RColorBrewer’ within R (http://www.r-project.org) and RStudio (https://www.rstudio.com). Differences in relative abundance between *V. parvula* alone and *V. parvula* + *P. gingivalis* groups were tested using LEfSe (*40*) considering 0.01 as the α value for statistical testing. These analyses included OTUs that were at least present in 20% of the samples.

## Acknowledgements

This work was supported by grants R21DE023967 (PID) and R01DE015254 (GH) and by the Intramural Program (NMM) of The National Institute of Dental and Craniofacial Research, National Institutes of Health. We also acknowledge the support of the Chilean National Fund for Scientific and Technologic Development (FONDECYT) grant 11180505 (LA).

## Author Contributions

PID, NMM, PDM and GH contributed to study design and supervised research. AH, HW and AM performed experiments. PID, AH, LA and BYH analyzed data. PID and AH drafted the manuscript. All authors read, critically revised and approved the manuscript.

