## Supplemental Figures and Table for "A cross-species interaction with a symbiotic commensal enables cell-density-dependent growth and in vivo virulence of an oral pathogen"

**Supplementary Material**

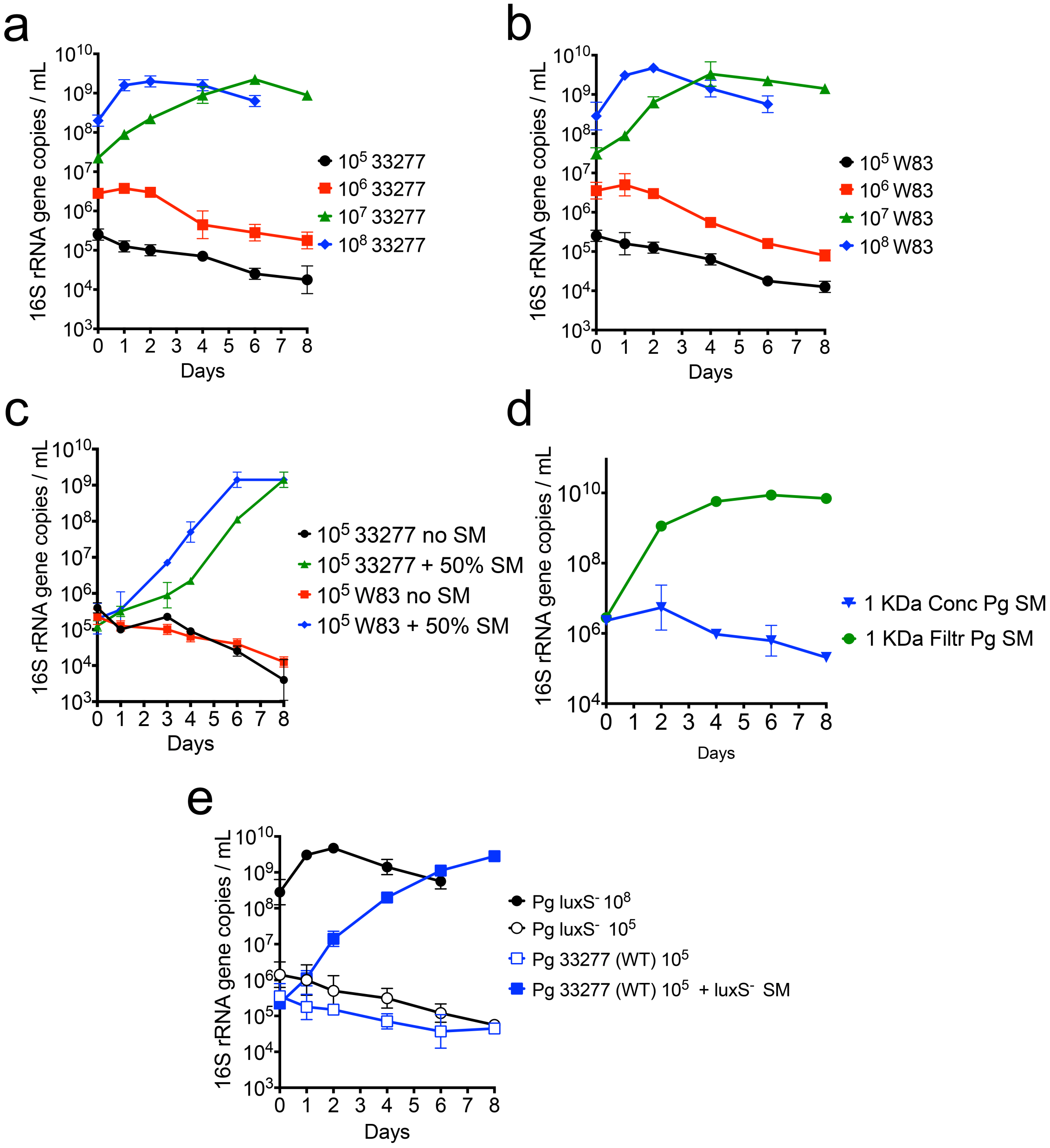


**Supplemental Figure 1. Cell-density dependent growth and effect of spent medium (SM) on different strains of *P. gingivalis* (Pg).** Growth of Pg ATCC 33277 (**a**) and Pg W83 (**b**) at different cell densities in mucin-serum. **c.** Effect of SM on the growth of Pg ATCC 33277 and Pg W83 in mucin-serum. **c.** Pg 381 soluble factor capable of supporting growth of low-cell-density Pg is smaller than 1 kDa. SM was obtained from

Pg 381 cultures grown in mucin-serum, then filtered through 3 kDa and 1 kDa membranes, followed by lyophilization and reconstitution (10x) in $dIH_20$. Reconstituted filtrates were added to fresh mucin-serum medium (1:3, vol:vol) to evaluate growth of low-cell-density Pg ($10^5$ cells $mL^{-1}$). **e.** Growth of a Pg Δ*luxS*::*ermF* mutant ($luxS^-$) at different cell densities in mucin-serum and effect of SM from the *luxS*$^-$ strain on growth of Pg ATCC 33277 (wild type).

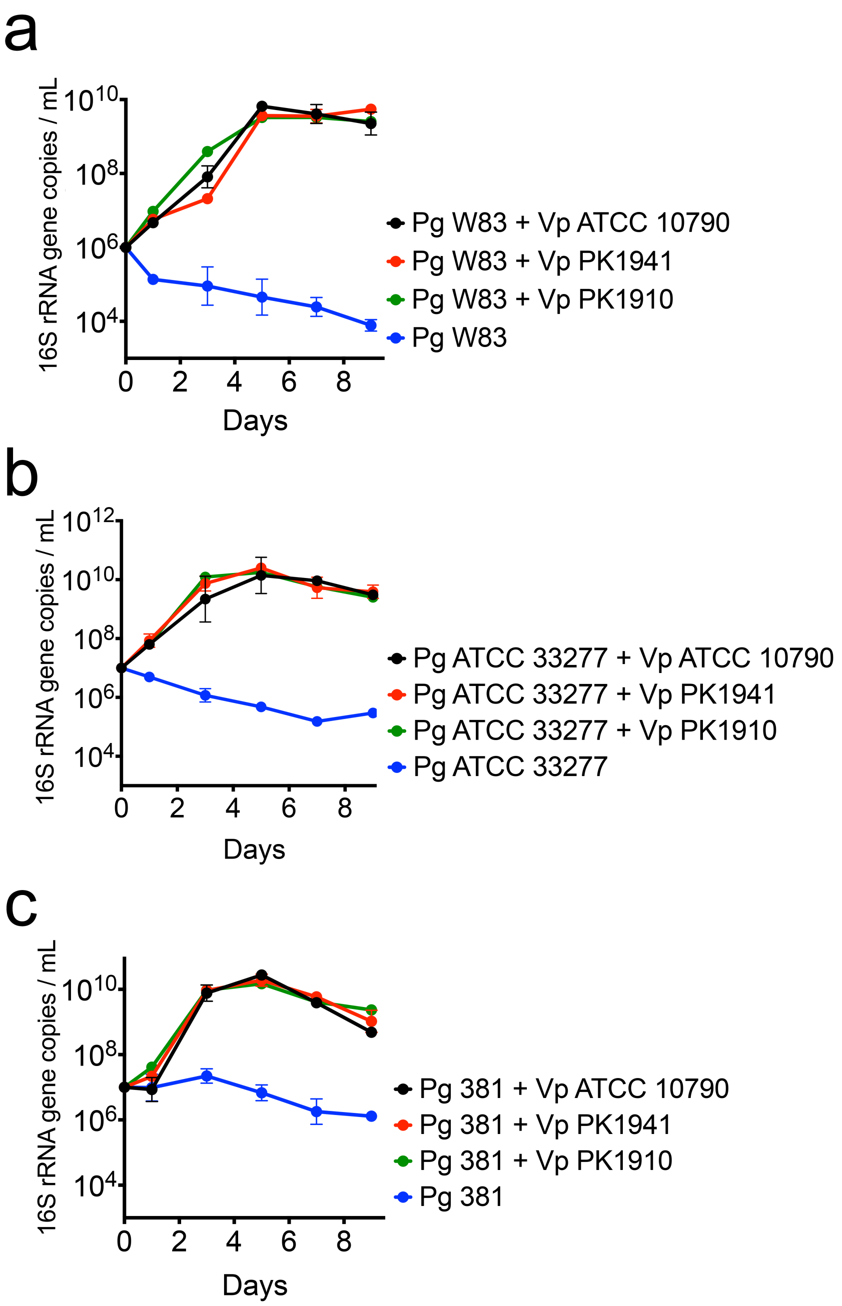


**Supplemental Figure 2. The interaction between *V. parvula* (Vp) and *P. gingivalis* (Pg) is not strain-specific.** Graphs show growth of Pg W83 (**a**), Pg ATCC 33277 (**b**) or Pg 381 (**c**) as monocultures and in the presence of either Vp ATCC 10790, PK 1941 or PK 1910. Inoculum size was $10^5$ cells mL$^{-1}$and all experiments were conducted in mucin-serum medium. Data represent mean and standard deviations from three independent experiments.

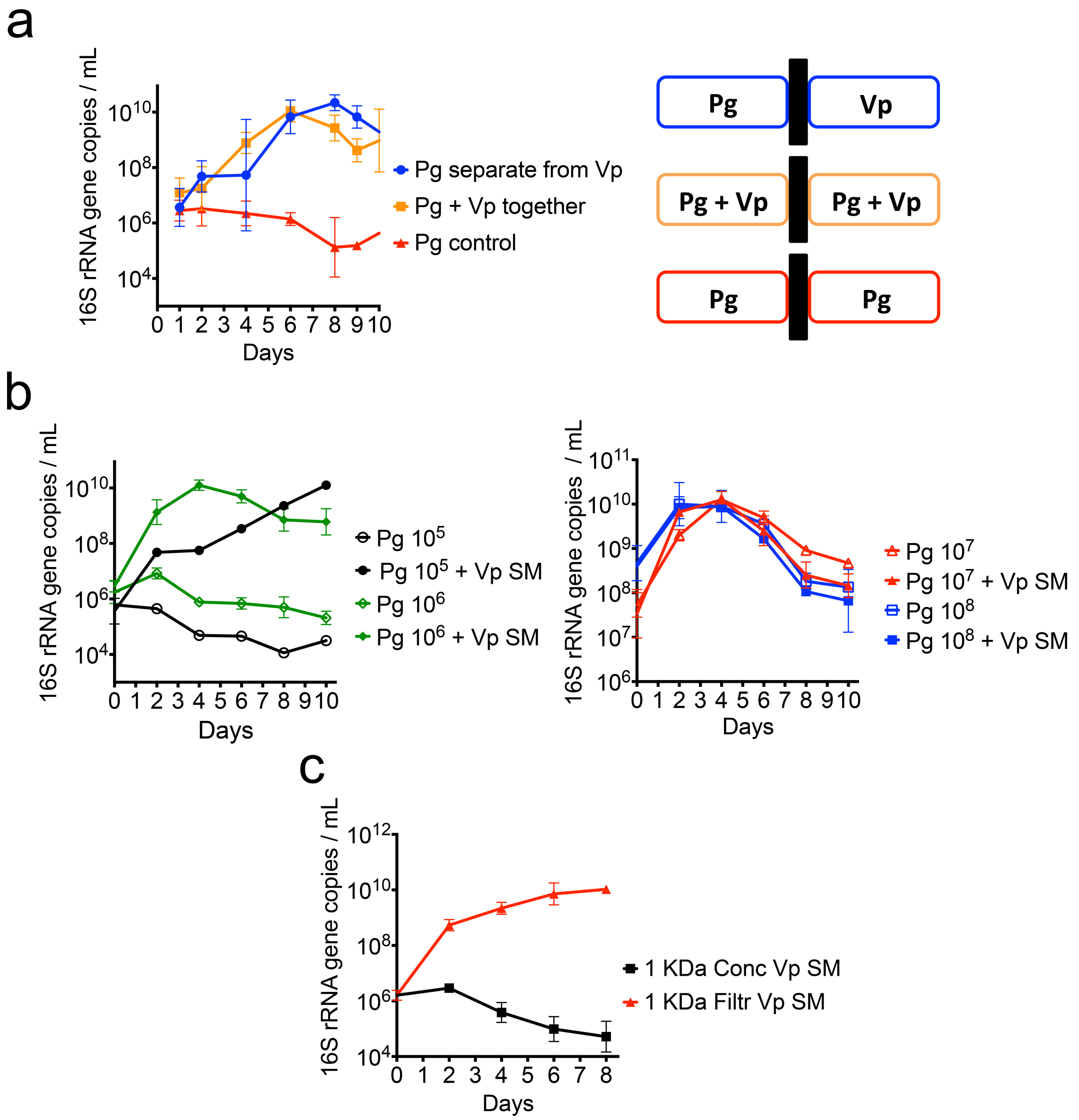


**Supplemental Figure 3. Characterization of the soluble factor mediating the support of *V. parvula* (Vp) on the growth of *P. gingivalis* (Pg). a.** Physical contact between Pg and Vp is not necessary to allow growth of a low-cell-density Pg inoculum. Pg and Vp were grown in chambers separated by a filter membrane with a pore-size of 0.22 µm. As controls, Pg + Vp were inoculated together or Pg was inoculated in monoculture at both sides of the membrane. Inoculation-density was $10^5$ cells $mL^{-1}$ for both species. **b.** Effect of Vp SM (24 h) on different Pg inoculum sizes. Left graph shows effect of Vp SM on Pg inoculated at $10^5$ or $10^6$ cells mL$^{-1}$, and right graph shows effect of Vp SM on Pg inoculated at $10^7$ or $10^8$ cells mL$^{-1}$. **c.** Vp soluble factor capable of supporting growth of low-cell-density Pg is smaller than 1 kDa. SM from Vp grown in mucin-serum was filtered through 3 kDa and then 1 kDa membranes, followed by lyophilization and reconstitution (10X) in dIH$_2$0. Reconstituted fractions (Conc= fraction >1kDa and Filtr=fraction < 1kDa) were added to fresh mucin-serum medium (1:3, vol:vol) to evaluate growth of low-cell-density Pg ($10^5$ cells mL$^{-1}$). Data in all panels represent replicates (mean and standard deviation) from at least three independent experiments.

**Supplementary Table 1.** List of compounds tested for their ability to induce growth of a low-cell-density inoculum of *P. gingivalis*. Experiments were performed with commercially available compounds in the indicated concentrations (middle column).

| Compound | Tested concentration (mM) | Effect on *P. gingivalis* |
|---|---|---|
| D-pantothenic acid (D-PA) | 0.005, 0.05 and 0.50 | No growth |
| D-panthenol | 0.005, 0.05 and 0.50 | No growth |
| β-alanine | 0.01, 0.10 and 1.0 | No growth |
| tyrosol | 0.002, 0.02 and 0.20 | No growth |
| spermidine | 0.069, 0.69 and 6.90 | No growth |
| spermine | 0.000049, 0.00049 and 0.0049 | No growth |
| cadaverine | 0.0979, 0.979 and 9.79 | No growth |
| putrescine | 0.1134, 1.134 and 11.34 | No growth |
| 4-aminobenzoate/para-amino benzoic acid (pABA) | 0.0729, 0.729 and 7.29 | No growth |
